# Crippled Coronavirus: 5’-PolyU targeted Oligo prevents development of infectious Virions

**DOI:** 10.1101/2022.03.04.483076

**Authors:** Hemayet Ullah, Saadyah Averick, Qiyi Tang

**Author notes:** Corresponding author: Hemayet Ullah.

## Abstract

Current RNA viral pandemic of COVID-19 has been worsened by rapidly spreading viral variants. To inhibit mutation-based development of new escape variants, elements that are indispensable for the virus may be targeted. The 5’-polyU tract of the antigenome offers one such target. Host cells do not harbor 5’-polyU tracts on any of their transcripts, making the tract an attractive virus-specific target. We hypothesize that inhibiting the 5’-polyU by complementary oligonucleotide can limit the use of the tract as template for virus to generate 3’ polyA tails of RNA viruses. Here, we used a frameshift-inducing DNA oligonucleotide with 3’ polyAs to target the 5’-polyU tract of mouse coronavirus (MHV-A59). Results from assays for double stranded RNA (dsRNA) synthesis, infectivity of released virions, and syncytium formation indicate that the oligonucleotide treatment prevented generation of infectious virions. Our results show a unique mode of action of the designed 3’-polyA oligonucleotide against mouse coronavirus which leaves host cells unaffected. This strategy can be adopted for the development of novel classes of oligonucleotide-based drugs that inhibit the production of infectious RNA viruses, including the coronaviruses. Since the 5’-polyU tract is conserved and is essential for variants of coronaviruses, this strategy can potentially address coronavirus variant emergence as well.

## Introduction

RNA viruses show increased capabilities for interspecies transmission, placing these viruses as the primary etiological agents of human-emerging pathogens, accounting for 44% of all infectious diseases [1]. These viruses use RNA dependent RNA polymerase (RdRp) for their genome replication and transcription. Since RdRp is error-prone, these viruses display high mutation rates in their genome, often allowing for the evolutionary adaptation of traits considered beneficial for viruses under changing environments [2–5]. The evolution and unprecedented rapid spread of the SARS-CoV-2 Omicron variant points to such capability inherent in many RNA viruses [6]. In addition to high mutation rates, recombination and reassortment processes have been attributed to the high biological diversity within the RNA viruses [3]. Their rapid adaptation to changing environments and increased biological diversity have proven difficult to overcome, and developing effective counter measures to these emergent new variants has proven challenging. Considering the current COVID-19 and past pandemics, it is quite plausible that by virtue of their mutational adaptability, RNA viruses will be the most likely candidates as the etiological agents of future pandemics. Though only a few drugs against RNA viruses have been developed [7], concerns also surround about the drug-escape-variants, like immune-escape-variants, which can evolve against any high efficacy drug that targets the virus. Given this, an effective strategy to develop durable and escape-proof countermeasures would be to target an area of the virus genome that is indispensable for the virus life cycle. Any mutation in the targeted indispensable part of the genome will not be tolerated by the virus; thereby, such an inhibition would limit the virus’s ability to mutate into a new variant in the presence of selection pressure from the inhibitor. In this regard, the 5’ polyuridines (5’-polyU) tract on the antigenome of RNA viruses which is used as template to generate 3’-polyA tail on their RNAs offers such a target.^1^

Many of the RNA viruses are cytoplasmic, and thereby cannot hijack the nuclear polyadenylation machinery to add 3’-polyA tails on their RNAs. However, it is now established that the 5’-polyU tract on the antigenome is used as a template to generate the 3’-polyA tail in the viral RNA genome and in the sub-genomic mRNAs while the poly-A tail of the genome serves as a template to generate the 5’-polyU tract on the antigenome [8–10]. Many positive-strand RNA viruses (e.g., members of the *Picornavirales, Nidovirales, Togaviridae, Caliciviriridae*, and *Astroviridae*) have long Poly(A) sequences at the 3’ termini of their RNA genome [8]. Minus RNA viruses use stuttering mechanism where RdRp moves back and forth over a stretch of 5-7 poly U residues at the 5’ end of RNA template and reiteratively copy the stretch into poly-A tail at the 3’ end of the viral mRNA [11]. Any mutation within the 5’-polyU tract will most likely prevent the virus from generating its 3’-polyA tail, which would effectively hamper the generation of its full-length genome to complete its life cycle. The essentiality of the homopolymorphic 5’-polyU tract on the antigenome for the survival and infectivity of positive RNA viruses, including many coronaviruses, is well-established [8, 12, 13]. Host cells do not harbor 5’-polyU tracts on any of their transcripts, making the tract an attractive, virus-specific target. Eukaryotic RNA polymerase III transcribes some small RNAs (t-RNA and 5S rRNA) with the 3’-polyU tract as a part of the transcription termination mechanism [14]. Molecular distinction between the viruses’ 5’-PolyU tract from the host’s 3’-polyU tract on transcripts is a central consideration to avoid any off-target effects.

Given that 5’-PolyU tract is essential for RNA viruses’ life cycle, we tested this hypothesis by using a molecular approach to target the 5’-polyU of a mouse coronavirus, mouse hepatitis virus (MHV-A59) antigenome. For the target, we used a complementary 3’-polydA-containing oligonucleotide. Use of the modified oligonucleotide against a fluorescent mouse coronavirus prevented release of infectious virions from the virus-infected cells. Application of the oligonucleotide successfully prevented the generation of fully formed vital replicase-transcriptase complexes (RTCs) as detected by the resident dsRNA specific antibody. This work represents a novel treatment strategy for many RNA viruses including coronaviruses, without negatively impacting host cells.

## Results

### Design of DNA oligonucleotide complementary to MHV-A59 5’ end of the antigenome (minus RNA genome)

Using the whole genome of sequences of MHV-A59 (accession# NC_001846), we carefully designed two oligonucleotides that were complementary to the polyUs at the 5’ end of the minus strand of the virus genome (Fig 1A). Considering that all the reported 21 PolyAs are used to make the 21 polyUs, the oligonucleotides were designed with twenty one 3’-polyAs and an additional 11 bases complementary to the flanking polyAs so that the designed oligos would be able to distinguish the 5’-polyUs from the 3’-polyUs. Also, the possible use of the oligonucleotide as a primer to amplify flanking sequence led us to make a frameshift mutation by deleting a base in the bison oligo-2 3’UTR flaking sequence. Additional modifications were done to the oligonucleotides to provide increased resistance to nuclease degradation, reduced toxicity, and to provide increased affinity for binding to complementary RNA (Fig. 1B). Though these kinds of modification have been used in antisense oligonucleotide technology to force usage of alternative polyadenylation site selection and to increase the abundance of message [15], here the use of a poly-A oligonucleotide, rather than a gene sequence-targeted oligonucleotide, can minimize any such effect on the genome. Though there are 11 flanking bases running into the 3’ UTR next to the N gene sequence of the virus, lack of any cryptic polyadenylation signal in the vicinity of the targeted sequence reduces any possibility of the use of an alternative polyadenylation site. Fig.1C shows the molecular structures of block copolymer-based oligo carrier nanoparticles which are described in detail below.

**Figure 1:**
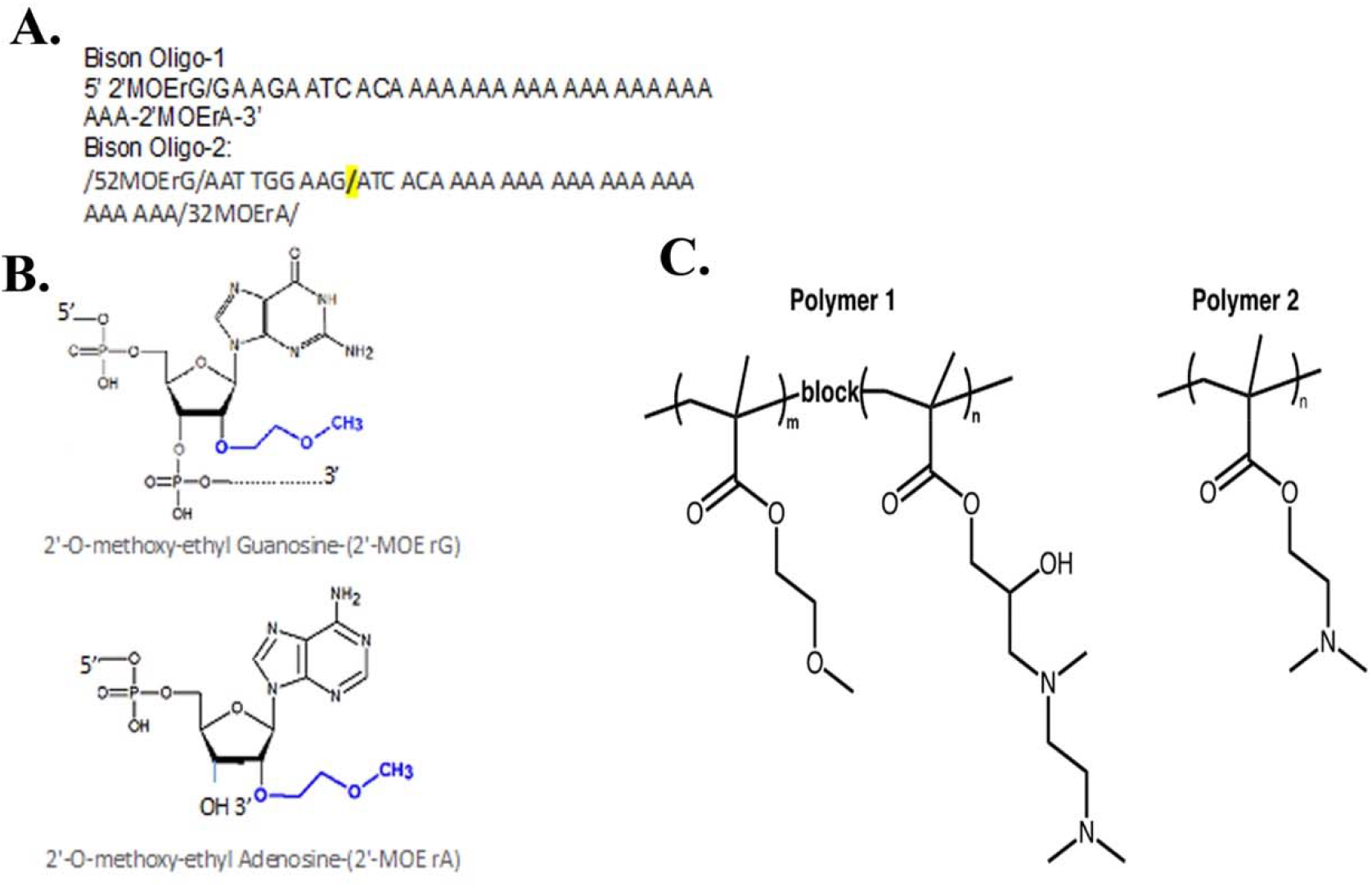
Sequence, modifications of DNA oligonucleotides used to target 5’-polyU of mouse coronavirus minus RNA. A. The oligo sequence is designed to be complementary to the 5’ end of the minus strand of Murine hepatitis virus strain MHV-A59 genome (Accession number NC_001846 of whole genome is used to generate the complementary sequence). Note 11 bases complementary to sequences downstream of 5-polyU is added to allow hybridization to the 5’-polyU instead of any cellular 3’-polyU tags. The slash shows the one base deletion from the 3’UTR sequence of the Bison Oligo 2. B. 5’ and 3’ modifications are (2’-O-methoxy-ethyl bases). C. Molecular structures of the block-copolymer (Polymer 1 and Polymer 2) used to deliver the oligonucleotides to the host cell.

### Oligo carrier nano-particles

Oligonucleotides, due to their anionic charge, size and nuclease sensitivity, have minimal cellular penetration, rapid elimination half-life, and can also induce an immunological response. To overcome these challenges of oligonucleotide delivery we used cationic polymers to complex and transfect the oligonucleotides to the infected cells.

#### Oligonucleotide delivery

To determine if the delivery vehicle would impact the efficacy of the Gen 2 DNA oligonucleotide, the oligo was formulated with either a lipid-based delivery agent (lipofectamine) or a polymer-based system. The oligonucleotide was complexed via electrostatic interactions of the negatively-charged DNA backbone and cationic charge on the lipid/polymer systems. The lipid-based oligonucleotide delivery systems are currently employed in the mRNA-based delivery of SARS-Cov2 vaccines, albeit with unknown lipid structures. The Polymer1 and Polymer2 system, are is a block copolymer with a first block that provides a hydrophilic biocompatible shell of oligoethylene oxide methacrylate and second block comprised of a hydrophilic tertiary amine methacrylate, polymer2 is a linear poly (dimethylamino methacrylate) and were prepared as previously described [16]. The polymer-based delivery systems were found to provide superior transfection (data not shown) and were used in subsequent experiments.

### Oligonucleotide-induced inhibition of infectious virion development

Mouse hepatitis virus (MHV) serves as a model for the family of enveloped plus RNA viruses from Coronaviridae and the MHV-A59 strain is the prototype coronavirus (CoV). We used the strain MHV-A59 harboring an eGFP fluorescent tag that was inserted by replacing a pseudogene-ORF4 [17]. Serial dilution of the stock virus was used to infect the mouse fibroblast cell line 17CL-1, which was derived by spontaneous transformation of 3T3 cells. As the designed oligonucleotides are expected to bind to the 5’-polyU site on the negative strand which develops after infection, we expected that the primary infection would not be affected by the oligonucleotide. Instead, we hypothesized that the oligonucleotide would interfere with the development of the +RNA genome and the sub-genomic mRNAs, which are synthesized in specialized Replication-Transcription Centers (RTCs). The primary infection-based development of infectious virions, which are released from the infected cells into the media, can be assayed to investigate whether the oligonucleotide interfere with the development of infectious virions. In this regard, we added NIR680 labelled secondary cells (magenta cells) to the primary cells 24h post infection and assayed for any virus-induced infectious effects in the secondary cells (Fig. 2). In control cells that were treated with the polymer alone, clear infectiousness of the virus was observed in the secondary magenta colored cells, which were mostly clustered in specific foci (Fig. 2A). Additionally, extensive cell-to-cell fusions to form syncytium- another indicator of infectious virion production were evident (arrow, Fig. 2A). On the other hand, within the oligonucleotide treated cells, only a few of the GFP-positive secondary magenta colored cells were seen and hardly any syncytium formation was seen within the secondary cells (Fig. 2B, 2C). In a few instances, some secondary cells in the presence of oligonucleotide did show GFP positive cells but as no cytopathic effects or any syncytium were evident in those GFP positive cells (data not shown), thus suggesting that the GFP positive cells in the oligonucleotide treated cells might not be producing fully formed functional virions.

**Figure 2:**
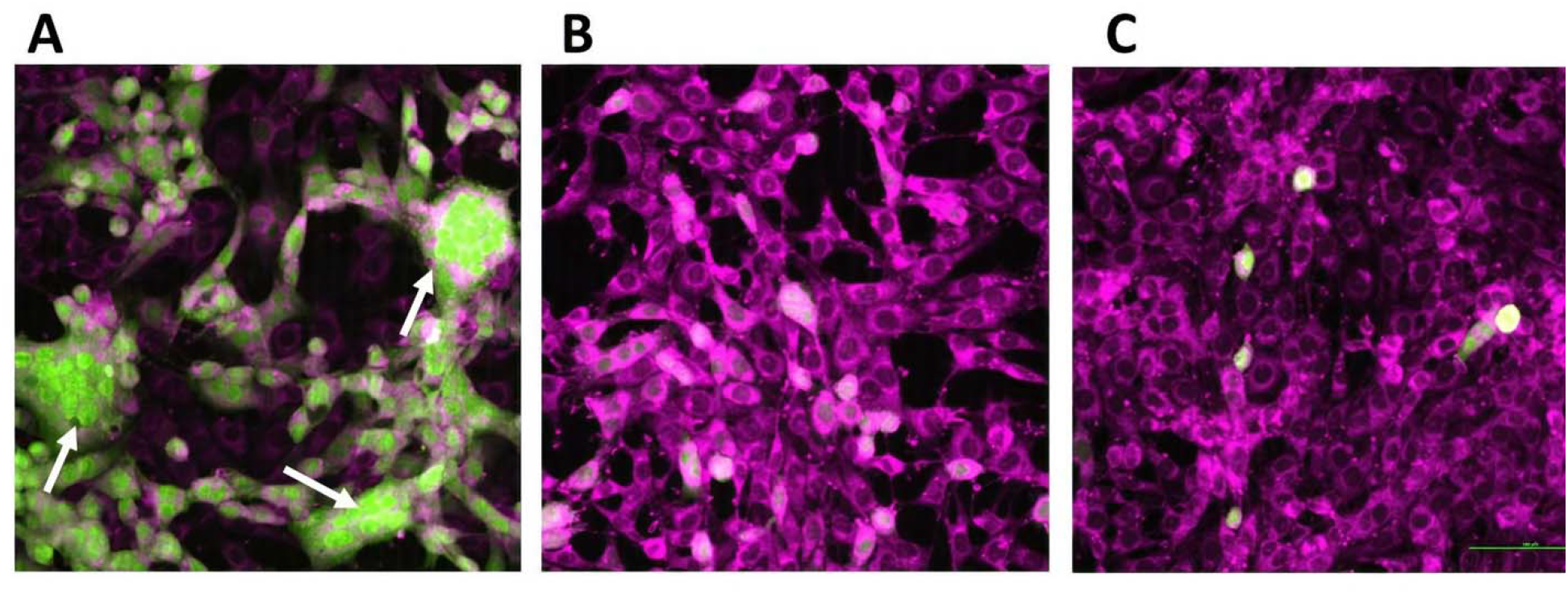
The oligonucleotides prevent MHV-A59-eGFP induced infection in fluorescently tagged secondary 17CL-1 cells (magenta colored). After 24h infection of primary cells, magenta colored secondary cells were added and imaged after 24h for infection of secondary cells. Image represents primary infection with virus at 0.1 MOI for 1h adsorption. A. MHV-A59-eGFP with polymer1. B. MHV-A59-eGFP with Polymer1 and oligonucleotide 1. C. MHV-A59-eGFP with polymer1 and oligonucleotide 2. Rounded, aggregated, fused, and granulated cells rapidly detaching from the monolayer are shown in panel A. Arrow in panel indicates infection induced syncytium (Original magnification 10X). Scale bar 100 μm.

### Oligonucleotide treatment interferes with the release of nucleocapsid protein from the host cells

To investigate whether the oligonucleotides are preventing the generation of fully functional virions in cells depicted in Fig.2, we isolated virion proteins from the same control and treated cell wells (from around the coverslip periphery) and media from the respective wells. We used western assay to detect viral nucleocapsid protein N, which is the most abundant viral protein produced in CoV-infected cells [18]. The N protein is the only viral structural protein that is known to bind to the RTCs and is essential in the incorporation of the viral genome into a fully functional virus particle [19–22]. Fig. 3A and 3B show the results obtained from the released virions in the media (Fig. 3A) and from the virus infected cells treated with or without the oligonucleotide (Fig. 3B). While both the biological replicates of released viral proteins in the media treated only with polymer1 showed extensive production of N-proteins, the released virions from cells treated with the oligonucleotides produced a non-significant level of N-protein. As expected, in media obtained from cells that were not treated with virus, we did not detect any N-protein. While cells treated with the oligonucleotide show slightly lower levels of N-protein production, compared to that of cells treated with the polymer only (Fig. 3B), the absence of N-proteins in the released virions indicates that the oligonucleotide treatment was indeed able to prevent the formation of the key structural protein required for the formation of infectious virions. Interestingly, when the background resolution of the blot was significantly enhanced, it could be clearly seen that polymer1-treated lysates obtained from the released virions in the media produced higher molecular weight bands, while no such bands were in the oligonucleotide-treated lysates (Fig. S1).

**Figure 3:**
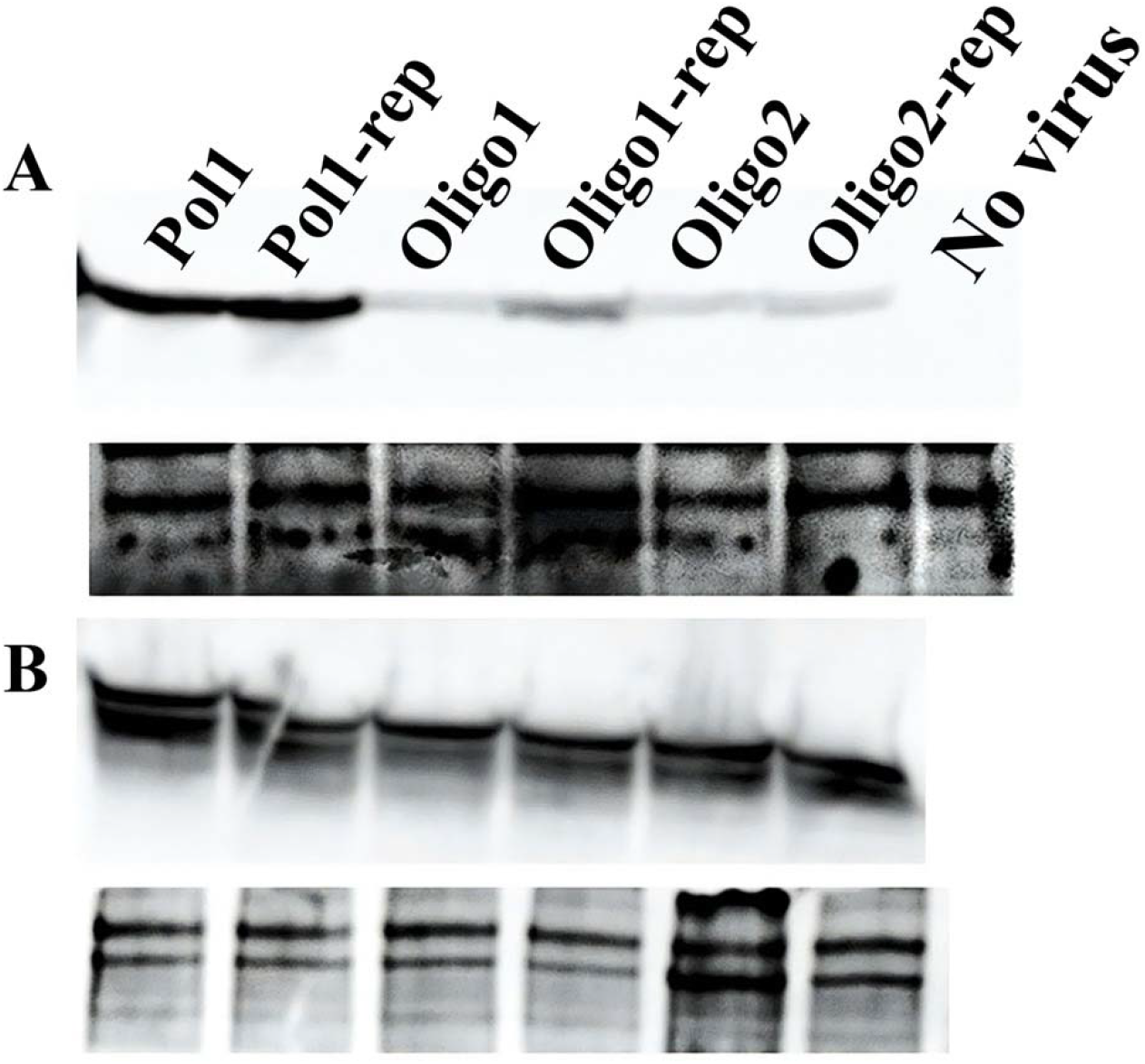
Detection of MHV-A59 nucleoprotein N in (A) released virions in the media (B) inside cell. In panel A, N-protein band of ~50 kD is shown in the virions from media collected from respective wells with cells shown in Fig. 2, freeze-thawed 3X. No virus treated media lysates is used as negative control. In panel B, lysates were collected from the respective wells of cells shown in Fig.2 (from around the periphery of glass slide). Respective treatments are shown on the top of each gel. As loading control, Coomassie stain of non-specific bands on the gel that was stained after the transfer of the majority of the proteins to the membrane are shown below each of the western band panels.

### Inhibition of Infectivity by virions obtained from oligo-treated cells

To confirm that the released virions from the oligonucleotide-treated cells are not infectious, we assayed the infection potential of the virions obtained from the media. The same media supernatants used in the western assay in Fig. 3 were serially diluted and used to infect the 17CL-1 cells for 24h and NIR680-labelled secondary cells were subsequently added. Because the cell lysates western showed almost similar levels of N-protein production across the control and treatments presumably from the primary infection (Fig. 3B), we wanted to evaluate whether similar levels of infectious virions were released in the medium. Therefore, without normalizing for the virus amount, we made serial dilutions of the medium supernatant and Fig. 4 shows the infection from the 10^-3^ dilution of medium containing the released virions from the respective treatment conditions. During the infection assay, no oligonucleotide or polymer was added. As can be seen in Fig 4, the 10^-3^ dilution of the media containing the released virions from the polymer-treated cells were successful in infecting the cells, as evident by the cytopathic effect as early as 6h post-addition of the secondary cells (Fig. 4 lower panel). The media from the oligonucleotide-treated cells, on the other hand, did not show cytopathic effects during the early infection phase. We also assayed for virion infectivity at 24h post-secondary cell addition. At 24h, the media containing virions released from polymer treated cells caused infection (GFP positive) in almost all of the magenta colored secondary cells (Fig. 4, upper panel). On the other hand, only few magenta colored secondary cells got infected when media containing virions released from the oligonucleotide-treated cells was used. the released virions media. Together, this infectivity assay confirms that treatment with the oligo prevented the release of infectious virions into the media.

**Figure 4:**
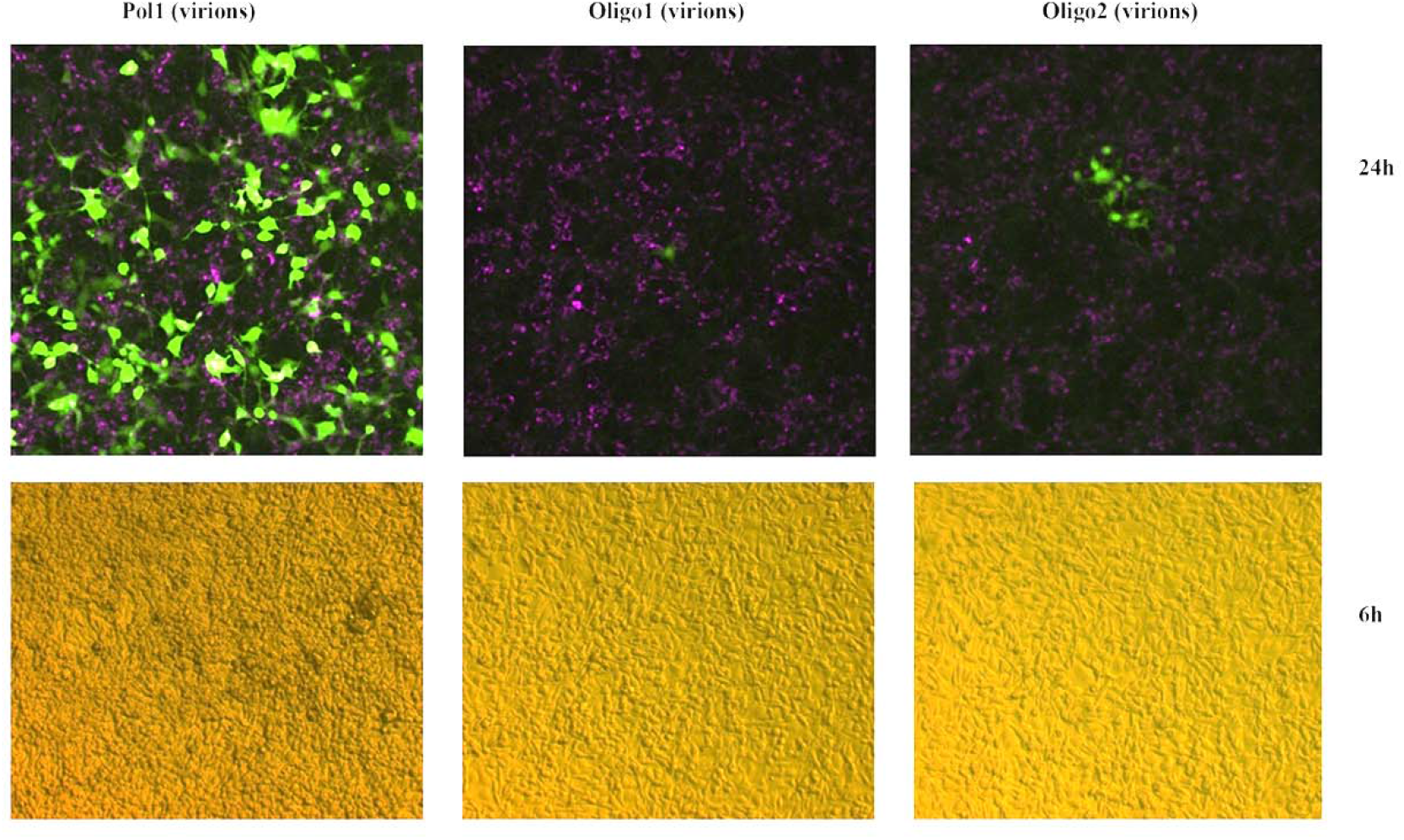
Infectivity assay done with the media released virions. Media collected from respective wells (infected with MHV-A59-eGFP +/− oligo) as depicted in Fig.2 were serially diluted and used to infect 17CL-1 cells for 1h, washed 3X times, then incubated for 24h in normal MEM media without any treatment of polymer and/or oligo. Media were replaced with NIR680 labelled 17CL-1 cells (0.25 x 10^6^ cells/ml) and were imaged after 6h (lower panel) in bright field and after 24h (Upper panel). The image shows the 10^-3^ dilution results in 10X magnification.

### Oligonucleotide inhibits development of double-stranded RNA intermediaries

We next assayed for RTCs in virus-infected cells to test for the presence of replicating virions in the oligonucleotide-treated conditions. In order to support viral replication and RNA synthesis, coronaviruses are known to develop double membrane vesicles (DMVs) by modifying cytoplasmic membranes in the perinuclear area to anchor RTCs [23]. During the active infection phase, coronaviruses produce various complete- and partially dsRNA intermediaries. These ds-RNA intermediaries are considered to be a marker for the location of active viral RNA synthesis in infected cells [24, 25]. Therefore, we set out to investigate whether MHV-A59 is able to establish active RNA synthesis processes in the presence of the oligonucleotide. As shown in Fig. 5, use of ds-RNA specific antibody indicated that the oligonucleotide significantly inhibited the development of ds-RNA intermediaries, while without the oligonucleotide, the virus actively developed replication/transcription intermediaries, suggesting active infection. Without the presence of active RNA synthesis mechanisms in a plus RNA virus such as MHV-A59, the formation of fully functional virions unlikely to occur. Therefore, it is likely that the oligonucleotide-induced inhibition of ds-RNA intermediary development is the underlying mechanism by which infection-incapable virions are produced in treated conditions.

**Figure 5:**
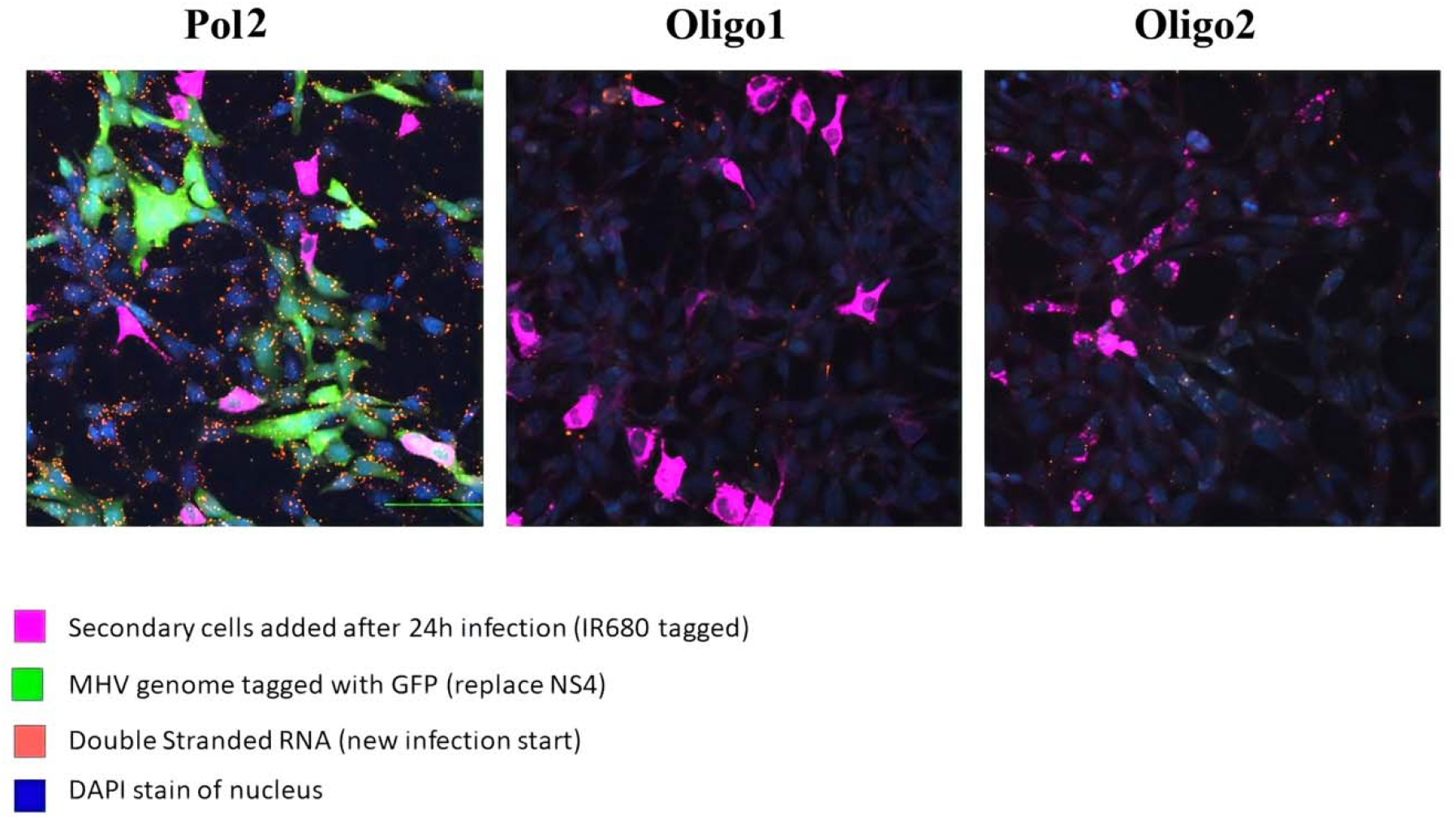
Immunofluorescence visualization of dsRNA during MHV-A59-eGFP active infection of 17CL-1 cells in the absence (panel A) or in the presence of oligo nucleotide 1 (B) or in the presence of oligonucleotide 2 (C). The cells were treated as described in the infection assays in the materials and methods section. The dsRNAs were observed by the TRITC labeled secondary antibody recognizing the anti-dsRNA primary antibody. The magenta colored cells were the NIR680 labelled secondary cells added to the primary cells infected with the MHV-A59-eGFP virus. Scale bar: 100 μm.

## Discussion

The less-than-perfect polymerase fidelity of RNA viruses drives successful adaptation to changing environments through the generation of genetic diversity. Even though RNA viruses with longer genome sizes – such as the coronaviruses – maintain greater polymerase fidelity by using exonuclease activity [26], the unprecedented capability of RNA viruses to cause and maintain pandemic-level threats is evident in the ongoing COVID19 pandemic. Targeting the virus in such a way that it becomes unable to generate escape variants via mutation represents a powerful way to contain RNA viruses. Here, we show that by targeting the viral indispensable 5’-polyU tract within the +RNA virus MHV-A59 [8, 12, 13], it is possible to inhibit the development of infectious virions. Given the indispensability of this tract, the likelihood of escape variant generation becomes low. Moreover, as host cells do not maintain any transcripts of their own with 5’-polyU tracts, the target offers a virus-specific, unique tag that can be targeted without affecting host cell transcripts. We show that the oligonucleotide-based approach is useful, as this strategy was successful in interfering with full virion assembly and release from infected cells. The oligonucleotide targeting the 5’-polyU prevented the ability of the virus to mount cytopathic effects through secondary cell infection. Though some leaky genome replication was observed, the prevention of infectious virion development can be used as evidence that the oligos binding to the polyU tract may prevent synthesis of fully functional virions. Further RNA binding experiments with the infectivity potential of the oligo-treated viruses will shed light on the underlying mechanisms driving the development of non-infectious virions. Here we have presented preliminary evidence establishing the 5’-polyU tag as a legitimate target for the successful containment of virus spread.

In a recent paper on the betacoronavirus mouse hepatitis virus strain A59 (MHV-A59) and on the alpha CoV enteropathogenic porcine epidemic diarrhea virus (PEDV), it was shown that virus endoribonuclease (EndoU) facilitates evasion of host pattern recognition receptor MDA5 [27]. The evasion happens by cleaving the 5’-polyuridines from the viral minus RNA strand which is a product of the 3’-polyA-templated RNA synthesis [27]. The paper suggested targeting the EndoU to allow the MDA5 to mount a robust interferon-based immune response. Though this result suggests that coronaviruses shorten the 5’-polyU tract to evade the host’s interferon-based immunity, shortened poly-U tracts are always found at the 5’ end of the minus RNA strand, indicating the essential need for the tract. In contrast to cellular mRNAs, in which the poly(A) is non-templated and is added post-transcriptionally, the viral poly(A) is template coded from a 5’-polyU tract in positive RNA viruses, such as in picornaviruses [28] and coronaviruses [24, 29]. Even though N-terminal RdRp-associated nucleotidyl-transferase activity can potentially add template-independent poly A, in HCoV-228E, a beta-coronavirus, it has been found that nsp8 (viral RdRp accessory subunit) has efficient 3’-terminal adenylyltransferase (TATase) activity only when the other strand has the 5’-polyU tract[30]. The oligo(U)-assisted/templated TATase activity on partial-duplex RNAs was confirmed for two other coronavirus nsp8 proteins, suggesting that the activity is conserved among coronaviruses [30]. The immature positive RNA strand becomes mature and stabilized only when iterative replication from the poly-U template adds the 3’-polyA tail [12]. It is quite possible that binding of the oligonucleotide to the 5’-polyU tag may mimic a shortened 5’-polyU stretch, allowing for short-term evasion of the host immunity response. This binding may prevent generation of fully functional and virulent virions with a full repertoire of infection capabilities. The inhibition of the all-important nucleocapsid structural protein by the oligonucleotide treatment clearly indicates that the virions produced would not be fully formed structurally, and thereby, would not be fully infectious. More mechanistic experiments will be able to establish the molecular details of such reduced capabilities to cause infection as reported in this study.

The study shows that the oligonucleotide was successful in preventing development of infectious virions by limiting genome replication activity, as evident by the inhibition of ds-RNA synthesis. Furthermore, the oligonucleotide successfully limited nucleocapsid protein synthesis in virions released from infected cells. The N-protein is known to form oligomers to constitute functional structures [31, 32], and N-protein self-association in the ribonucleoprotein complex is essential for producing fully functional virions[32]. Use of native gel-based western assays can be used in the future to investigate this effect of the oligonucleotide on the N-protein oligomerization. However, it is known that MHV-A59 structural proteins, such as nucleocapsid N-protein, are not required for viral RNA synthesis, as the virus replicase gene products suffice for discontinuous transcription [33]. It is quite possible that the role of coronavirus N-protein is to evade host immune-surveillance by antagonizing the interferon (IFN) response [34, 35]. It has also been reported that coronavirus N-protein can act as a suppressor of host-induced viral RNA silencing [36]. Interestingly, all identified suppressors of RNAi from mammalian viruses possess IFN antagonistic properties, suggesting that RNAi and innate antiviral responses are interrelated [36]. Both of the host antiviral responses-IFN and RNAi based, evasion by the N-protein requires the ds-RNA function [35, 36] and the oligonucleotide treatment in this study affected both the N-protein and ds-RNA synthesis significantly, indicating a potential mechanism of oligonucleotide function reported here. This simple approach of preventing potential genome replication represents a novel avenue to counter error-prone RdRp-based rapid evolution and adaptation of mutant variants of RNA viruses. Precise elucidation of the molecular effects of oligonucleotide-induced viral disruption will warrant future studies. Specifically, studies identifying the fate of the oligonucleotide-induced plus/minus genome with regards to stability, PolyA/U length, and replication potential should be conducted. In addition, the effect of the oligonucleotide on host cell ds-RNA sensors, interferon production, and immune evasion potential of the virus can be studied further.

The successful application of the oligonucleotide to target the mouse coronavirus 5’-polyU tag can be attributed to the unique design of the oligo, stability modifications, and to the use of block-polymers to deliver the oligos to the cell interior. The coronaviruses do not enter the host cell nucleus, where host cell mRNAs are processed by nuclear-localized enzyme; instead, coronaviruses are known to replicate in the cytoplasm [37]. This may allow the oligos to bind to their target easily. Binding of the oligonucleotide can easily affect the synthesis of the nucleocapsid protein as well. In addition to the 3’-polyA tail, all of the MHV A59 plus RNA strands that include the whole genome and 6 subgenomic mRNAs possess the 1.7-kb sequence of RNA-7 [25]. RNA-7 contains the N gene sequence next to the oligonucleotide binding site. Therefore, it is quite possible that binding of the oligonucleotide to the target site may interfere with copying the minus strand to generate the positive RNA strands with the N-gene sequence. Future in-depth transcriptomic analyses will be able to shed mechanistic insight into the oligonucleotide effect on the viral genome.

The above discussion indicates that the positive sense RNA viruses essentially require a 5’-polyU tract to complete their life cycle and to successfully infect host cells. Therefore, inhibiting/blocking the 5’-polyU tract should also interfere with this key survival mechanism of the virus and may lead to the production of a ‘dead’ genome or to a crippled virus being unable to infect. Targeting the 5’-polyU with lipid nanoparticle wrapped poly-A or with oligo dA, or DNA aptamer, or 5’-PolyU binding small compound and/or protein can limit the use of the minus strand for generating the positive sense RNA strand genome and the subgenomic mRNAs with polyA tails.

In addition, it has not escaped our attention that the poly-U tract is immediately generated upon infection [29]. As such, the tag can be leveraged and targeted for early detection of viral infection as well. This may overcome many of the issues encountered with the prevalently used real-time PCR-based detection systems, such as the lack of enough templates early in the infection process, which may drive false negative results.

Scientists around the world are working to understand this devastating illness and help put an end to the terrible toll caused by the COVID-19 pandemic. Experimental establishment of an indispensable target like this 5’-polyU tract on the antigenome of coronaviruses may assist in the development of effective drugs that prevent the generation of drug/immune-escaping virus variants.

## Acknowledgements

The following reagents were obtained from BEI Resources (NIAID/NIH): Recombinant Murine Coronavirus MHV-A59 with Enhanced Green Fluorescent Protein (eGFP) (Cat# NR-53716), Murine 17Cl-1 Cell Line (derived from 3T3 cells) (Cat# NR-53719), and monoclonal anti-murine coronavirus nucleocapsid (N) protein, Clone 1.16.1 (produced *in vitro*) (Cat# NR-45106). We thank Dr. Raphael Itay and Ms. Aniqa Tasnim for critically reading the manuscript. Immunofluorescence images were obtained with a spinning disk confocal fluorescent microscope acquired through a Department of Defense HBCU/MI Equipment/Instrumentation Grant (#64684-RT-REP) to Dr. Anna K. Allen of Howard Biology Department.

## Competing interests

The authors have no competing interests

## Data and material availability

all data is available in the manuscript or the supplementary materials.

## List of Supplementary Materials

### Supplementary Materials

Materials and Methods

Fig S1

## Supplementary Materials

### Materials and Methods

#### Cells and Viruses

Murine 17CL-1 cell line (derived from 3T3 cells) was obtained through BEI Resources, NIAID, NIH, catalog number: NR-53719. The cells were maintained as monolayer cultures in Minimum Essential Medium (MEM; Sigma Aldrich-M 4655) containing 10% fetal bovine serum (FBS; Life Technologies), 100 IU/ml of penicillin, and 100 μg/ml of streptomycin (Both from Life Technologies) in a 37°C humified incubator supplemented with 5% CO2. MHV strain A59-eGFP, which express the Enhanced Green Fluorescent Protein (eGFP) inserted in place of the Ns4 gene, was obtained through BEI Resources, NIAID, NIH, catalog number: NR-53716.

#### Infection Methods

The 70-80% confluent 17CL-1 cells were pretreated with the oligo (6 μg) plus polymer (1 μg oligo / μg of polymer) or polymer only for 1h at 37C inside 5% CO2 supplemented incubator. Cells were then infected with the MHV-A59-eGFP at 0.01 MOI. After one hour of adsorption, the media were removed and the cells were washed three times with PBS buffer. New MEM media with oligo plus polymer or polymer only was added to the virus treated cells and was incubated for 24h. Trypsinized 17CL-1 cells from non-treated wells were pelleted (10^6 cells / ml) and then labelled with diluting dye (BioTracker™ NIR680 Cytoplasmic Membrane Dye Live Cell Dye, Sigma-Aldrich, Cat. # SCT112) 1:2000 in culture medium for a final dye concentration of 1uM. After 20 minutes of incubation inside 37C cell culture incubator, cells were pelleted at 350xg for 5 minutes. The pelleted cells were washed 3 times with cell culture medium and 0.25 x 10^6 / ml resuspended secondary cells along with oligo plus polymer or polymer only added to the primary cells already infected with the virus and the cells were assayed at the indicated times of incubation.

#### Protein Extraction, Western assay

Before protein isolation, cells were washed three times with cold PBS. Lysates were isolated in lysis buffer (Cell Signaling, Danvers, MA) supplemented with protease inhibitor (Sigma-Aldrich), phosphatase inhibitor cocktail A and B (Santa Cruz Biotechnology, Dallas, TX), and N-Ethylamaleimide (Sigma-Aldrich, St. Louis, MO). The concentration of all proteins was measured using Bio-Rad’s Bradford dye following the manufacturer’s instructions. Lysates from media supernatants were prepared by three quick freeze-thaw cycles of the media. After 50 microgram of cell lysates or equal amount of supernatant lysates with 1X loading dye (Biorad, Hercules, CA) supplemented with 50 μl/ml βME (beta-mercaptoethanol) were boiled for 10 minutes, chilled lysates were loaded on the BioRad’s 4-12% precast Bis-Tris polyacrylamide gel, Proteins were then transferred onto a precut PVDF membrane (Biorad) using Biorad electrotransfer set up. After transfer, the gel was incubated overnight with Coomassie dye to stain the left-over proteins as loading control. The membrane was blocked with 5% nonfat milk in PBST (1X PBS with 0.1% Tween 20) for one hour, three washes were done for 10 minutes each by using PBST (1X PBS with 0.1% Tween 20). Washed membrane was incubated overnight at 4C with the indicated diluted primary antibody, followed by incubation with a horseradish peroxidase coupled secondary antibody. After washing three times with PBST, the protein expression was visualized by Biorad’s Clarity ECL reagent using Protein Simple Fluor-Chem M system.

#### Antibody, Immunofluorescence microscopy

The antibody directed against double-stranded RNA was obtained from UniQuest Pty Limited- the commercialization company of the University of Queensland, Queensland 4072, Australia, (Cat# MMABA 2G4). Monoclonal Anti-Murine Coronavirus Nucleocapsid (N) Protein, Clone 1.16.1 (Cat# NR-45106) was obtained from BEI Resources, NIAID, NIH.

#### Immunofluorescence Assay

Briefly, the cells on glass coverslip in 12 well plate were incubated with the treatments as described in the Infection method section. After the indicated time frame, the cells were washed 3X with PBS, fixed with 4% PFA-PBS for 15 minutes. The permeabilization was done at room temperature with 0.5% Triton X-100 in PBS (pH 7.4) for 10 min. Cells were washed 3X with ice cold PBS. For blocking, the cells were incubated with 5% BSA, 22.52 mg/ml glycine in PBST (PBS with 0.1% Tween 20) for 30 min at room temperature. The washed cells were incubated in a humified 4C chamber overnight with the anti-dsRNA antibody (1:250) in PBST with 1% BSA. The cells were washed three times in PBS, 5 min each wash and then Incubated cells with the anti-mouse TRITC conjugated secondary antibody (1:1000) in 1% BSA for 1 h at room temperature in the dark. After washing three times with PBST for 5 minutes each, the cells were mounted on slide using Antifade mounting media with DAPI (4-6-diamidino2-phenylindole; ThermoFisher). After sealing the coverslips with clear nail polish, the slides were left at RT overnight for curing. Imaging was performed using an Eclipse Ti 2000 laser-scanning confocal microscope (Nikon CSU series Spinning Disk confocal microscope) with DAPI, FITC, TRITC, and Cy5 filter sets.

#### Figure S1

**Fig. S1.**
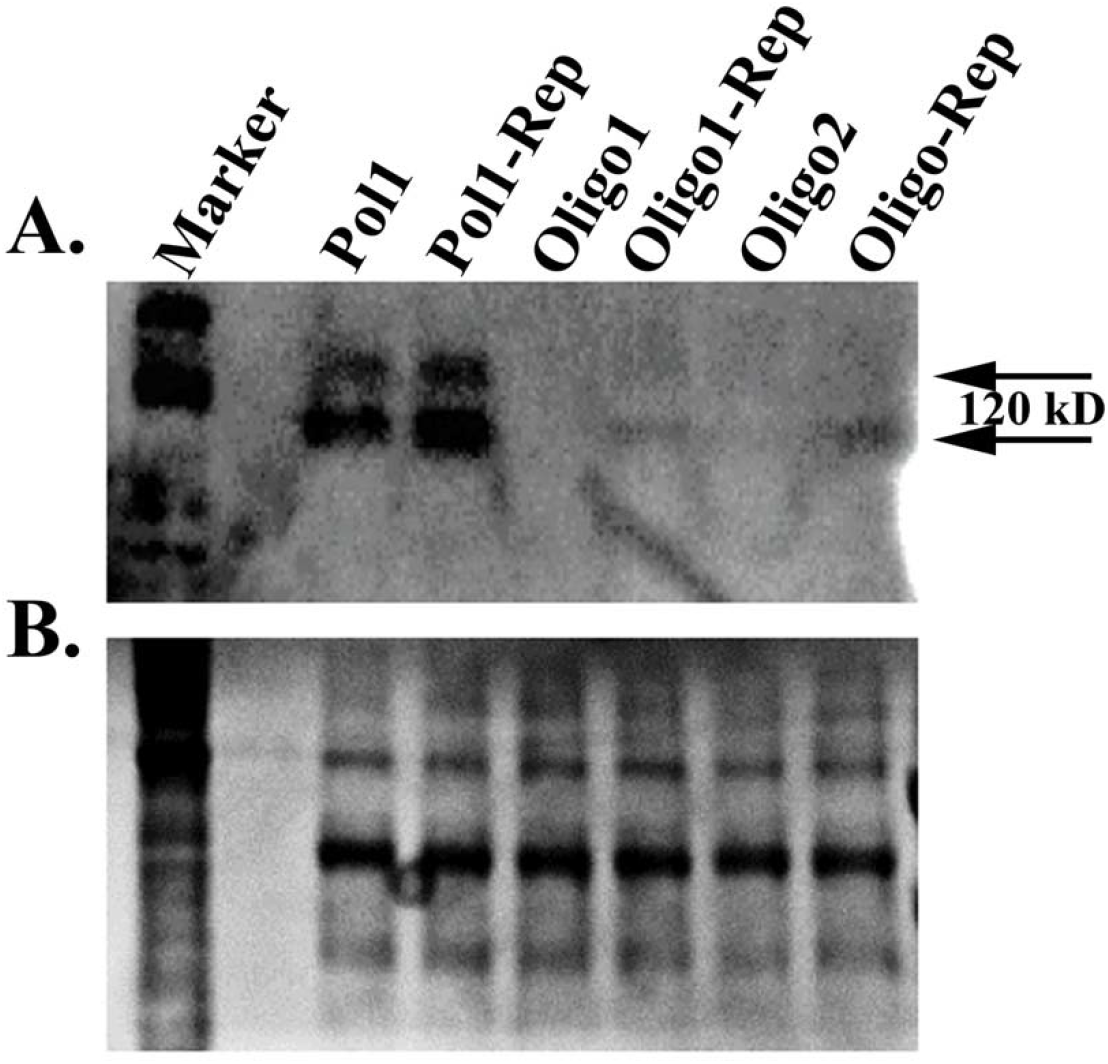
Enhanced background of western blot image revealed the potential higher molecular weight oligomeric bands of MHV-nucleocapsid (N) protein in the lysates obtained from the media used in Fig. 3 western assay indicated as lane Pol1 and Pol1-replicate. No such higher molecular weight bands could be seen in the lysates obtained from the media of the wells treated with oligo 1 or with oligo 2. As loading control, Coomassie stain of non-specific bands on the gel that was stained after the transfer of the majority of the proteins to the membrane are shown below the western band panels. Note that the bands were seen on a denaturing SDS-PAGE gel only after uniform enhancement of the blot background.

## Footnotes

1 Provisional patent application # 63025613, EFS ID# 39456605 (Title: Targeting the Unique minus Strand mRNA to Stop Replication of COVID-19 Causing SARS-CoV-2 virus without affecting host cells). PCT application submitted May 14, 2021.

## Notes

### Competing Interest Statement

The authors have declared no competing interest.

